# From Concrete to Canopy: Illuminating Moth Biodiversity in New York City’s Urban Jungle

**DOI:** 10.1101/2025.07.11.663976

**Authors:** Shira Linsk, Anna Thonis, Kristin M. Winchell

## Abstract

Moths (Lepidoptera) are sensitive to anthropogenic threats and serve as valuable bioindicators. Despite the remarkable diversity and abundance of Lepidoptera globally, there is a lack of information on how moth species are impacted by urbanization. Notably, very little is known about moths in the largest city of the United States, New York City, where pervasive urban pollutants, artificial light at night, land cover change, and habitat fragmentation are severe. We examined the effects of urbanization on moth biodiversity in New York City, with a focus on green spaces. We used citizen science records from *iNaturalist* and complemented these data with ground sampling at twelve locations across six parks at night. While the *iNaturalist* dataset is comprehensive both spatially and temporally, it failed to detect rarer species we observed on the ground. However, the scope of the field survey dataset is limited in geographical breadth and seasonal coverage. Overall, we found a negative relationship between greater urbanization and moth diversity, with community similarity correlated with environmental similarity. Our results found greater biodiversity with less light at night and urban development, and more deciduous tree cover and open land. Our structural equation model reveals additional insight: although we detected a strong direct negative effect of developed land on moth diversity, urbanization also negatively impacts diversity via indirect effects of reducing open space and deciduous tree cover. Developed open space alone does not directly affect diversity but may positively impact diversity through its covariance with vegetation cover. These findings support the importance of mitigating artificial light at night in urban green spaces and maintaining urban vegetation to ensure nocturnal Lepidoptera can persist in rapidly urbanizing landscapes.

## Introduction

Urbanization is linked to declines in biodiversity through biotic homogenization and species extirpation [1–2]. Although these impacts are seen across the tree of life, the effects of urbanization on invertebrates have been particularly drastic, with steep declines across insect orders [3–4]. Studies have demonstrated local extinctions of insect populations and reduced insect diversity with increasing urbanization [5–6]. Much of this work has focused on bees and butterflies, revealing decreased diversity and abundance with increased urbanization [5, 7–9]. Urbanization presents many challenges for insects, including the urban heat island effect, habitat fragmentation, pollution (including artificial light), impervious surfaces, and exotic plants [10]. Habitat fragmentation and landscape maintenance are major contributors to pollinator decline, including Lepidoptera [11]. However, alongside these challenges come opportunities in terms of anthropogenic resources that may be exploited, as seen in common urban species like cockroaches, flies, and carpet beetles [12]. Many species may thrive in cities despite the threats of pest control and habitat modification. For example, urban plantings may generate habitat and resources that support invertebrate diversity, including Hymenoptera and spiders [13–16].

Relatively little research on urban invertebrates has focused on moths, and there is a notable bias toward butterflies and other diurnal Lepidoptera [7, 17–18]. This is perhaps due to the challenges associated with observing moths at night (e.g., low visibility, general safety considerations) and the minimal public engagement surrounding them compared to butterflies [19]. Some recent studies on moths have found decreasing populations with increased artificial light at night [20–22]. Additionally, anthropogenic environments generate intense selection pressures and filtering processes that may favor the survival of only a few species, such as heat-tolerant moths [23] or produce phenotypic shifts, such as increased body size in urban moths [24]. Indeed, studies of industrial melanism in moths represent some of the first examples of rapid evolutionary change in cities [25–27].

Understanding the effects of urbanization on moths is important because of the crucial roles they play in ecosystem services (e.g., pollination, food, and nutrient cycling). Moths are fundamental to ecosystem health and stability and serve as bioindicators [22, 28–29] since they respond quickly to environmental change [20, 30–31]. There are approximately 150,000 known species of moths globally [32], and hundreds of new species are described annually [33]. Yet, we know very little about the species that persist in urban landscapes. For conservation initiatives to be successful in these spaces, we must first take stock of the species that reside there and relate patterns of diversity to urban stressors.

We investigate patterns of moth diversity in New York City, where artificial light at night, pollution, and habitat fragmentation are extreme. We hypothesize that landscape-scale features influence species diversity and predict that developed land use negatively impacts moth diversity. In addition, we hypothesize that habitat-scale features such as pollution, light at night, and microclimate correlate with moth diversity. By connecting moth diversity to urban habitat features, we aim to improve our understanding of the ecological impacts of urbanization on nocturnal Lepidoptera species in one of the largest urban centers globally.

## Methods

Statistical analyses were conducted using R studio (ver. 2023.12.0.369) [34] and R (ver. 4.3.2) [35]. For all linear models, we evaluated multicollinearity with the ‘VIF’ function in the package *car* [36] assessed statistical significance using t-tests for individual coefficients, and evaluated model fit based on Akaike Information Criterion (AIC) and residual deviance.

### iNaturalist data

We downloaded 31,793 Research-Grade observations of the Order Lepidoptera using a polygon spatial filter around all five New York City boroughs. We manually removed butterfly Families (i.e., Papilionidae, Nymphalidae, Pieridae, Lycaenidae, Hesperiidae), resulting in 12,956 moth observations. We imported records into QGIS (ver. 3.34.3-Prizren) [37] and used the New York City administrative boundary layer [38] to reduce this dataset to 10,740 records located within New York City boundaries. We used the plugin *Density Analysis* [39] to quantify observation density using a grid of 1 km hexagonal cells across the region (Fig. 1). We excluded hexagonal zones (33 zones) with undefined landscape types or >70% water cover (according to the National Land Cover Database “NLCD” [40]), to ensure the zones had relevant habitat data for terrestrial species. We filtered the *iNaturalist* records to include observations from 2000 onwards to better capture moth diversity over time, rather than at one specific point in time. Additionally, we removed moth observations that we ourselves contributed. This resulted in a total of 9,374 records.

**Figure 1.**
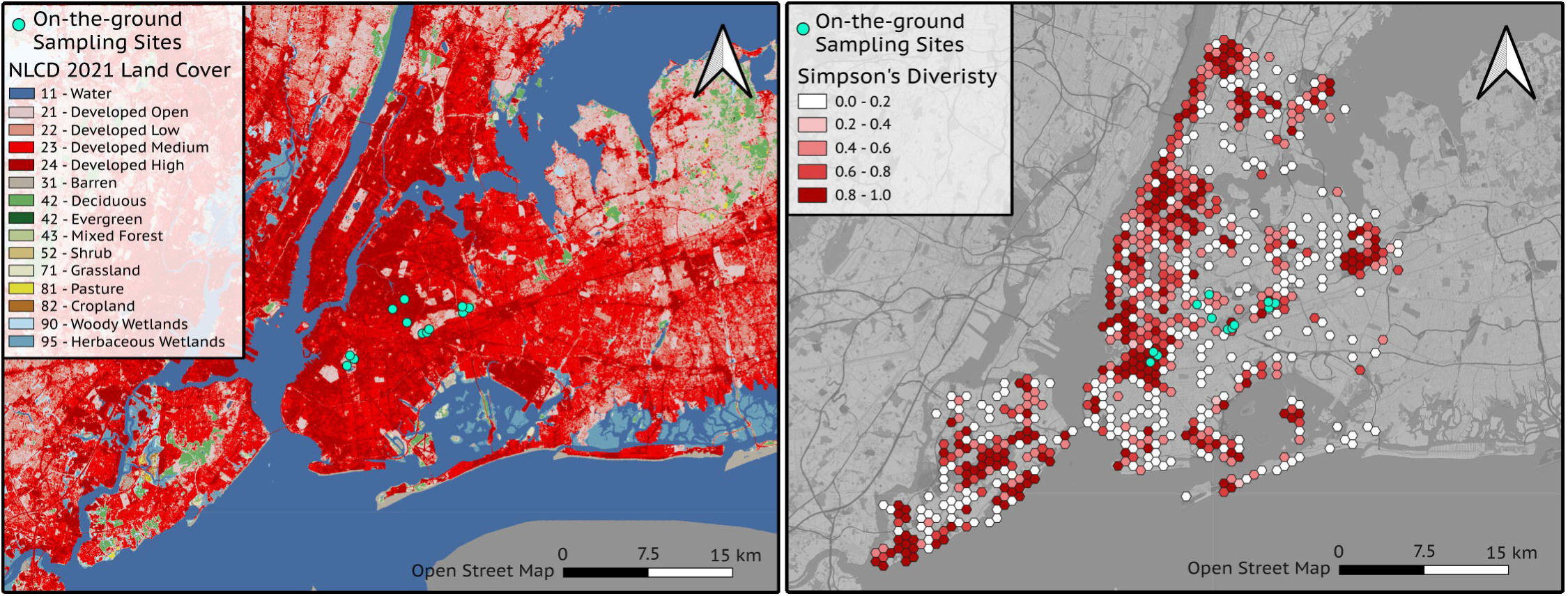
Study sites and moth diversity patterns across New York City (a) On-the-ground sampling locations for 12 locations in Brooklyn and Queens, New York City, USA, where we sampled Lepidoptera, and National Land Cover (NLCD) data for New York City. (b) Species diversity in 1-km hexagonal bins derived from iNaturalist data. Base maps by Open Street Map.

### Field Site Selection

To complement our *iNaturalist* dataset and provide field-based comparisons with any observed trends, we identified 12 on-the-ground sampling sites in six greenspaces across Brooklyn and Queens, New York, that reflect variation in urban-associated land cover and habitat features encountered across the city (Fig. 1). We visually assessed satellite imagery to identify green spaces and then chose sampling locations within each space spanning a gradient in light and air pollution based on on-the-ground measurements (rather than remote-sensed) to capture variation in this variable at an organismally relevant spatial scale.

### Field Sampling

We surveyed each location for one hour shortly after dusk (8-10 pm) between March 3-April 29, 2024. The exact time at which sampling occurred varied throughout our survey timeframe as sundown began later into the evening as the season progressed. We restricted the temporal sampling period to minimize seasonal variation in species detected across sites. We constructed a light trap using a suspended 5 m^2^ white sheet directly illuminated with two flood lights (6000 Kelvin, visible light spectrum) and one ultraviolet LED light (385-395 nm) (as in [41]). Moths are easily sampled with light traps, allowing for robust estimates of species richness and abundance [30]. Each survey consisted of two people photographing all moths observed landing on or in the immediate vicinity (on illuminated ground within 2m) of the light trap. We identified species from the photographs and counted the total number of individuals per species encountered during each sampling event. In addition, for each sampling event, we measured the following site-level variables: air pollution (PM 2.5; Temtop M2000C), light at night (lux; EXTECH Light Meter LT300), temperature, humidity, and wind speed (Kestrel 3000).

### Landscape analysis

We mapped *iNaturalist* records and on-the-ground sampling locations using QGIS. We extracted zonal statistics of NLCD land cover classifications [40] within each 1 km hexagonal zone for the *iNaturalist* dataset. For on-the-ground sampling locations, we used a 150 m buffer around each sampling point to capture the immediate landscape characteristics of each site and at a spatial scale relevant to Lepidoptera community structure (100-200 m) [42]. We calculated the percentage of each land cover class for hexagonal zones and buffered areas. We summarized percentage of land cover classifications (i.e., Deciduous, Mixed Forest, Grassland, Pasture/Hay, Developed Open Space, Developed Low-Intensity, Developed Medium-Intensity, Developed High-Intensity) with principal component analysis (PCA) using the function ‘prcomp’ in base R *stats* [35]. We visualized results with packages *factoextra* [43] and *corrplot* [44].

### Statistical analyses

We summarized species diversity with Simpson’s Diversity Index (Simpson’s D), which considers both species richness and evenness [45]. Simpson’s Diversity Index is robust in urban settings as it is not sensitive to rare taxa that may be under-sampled or absent in disturbed habitats where common generalist species might dominate [23]. Additionally, this index can overcome issues of small sample size and spatial biases [46]. We calculated Simpson’s D (1-D) using species-level identification for each *iNaturalist* hex bin and each on-the-ground sampling location in R using the function ‘diversity’ in the package *vegan* [47].

We conducted three analyses using the *iNaturalist* data. First, we used multiple matrix regression (MRM) to analyze the similarity of community composition with respect to environmental similarity across sites This entailed generating a distance matrix (Euclidean Distance) of species composition similarity across hexagonal zones, as well as for the environment (percentages of all NLCD land cover classes). We used the function ‘dist’ in the R base *stats* for all distance matrix analyses [35] and performed matrix regression using the function ‘MRM’ in R package *ecodist* [48]. Second, we used linear regression to analyze the relationship between moth diversity (Simpson’s D) and landscape-level (remotely sensed) habitat features. Explanatory variables in the model included NLCD categories (as a proportion of hex-bin area): deciduous tree cover, developed land cover (high-, medium-, low-intensity), pasture, developed open land, shrub, and grassland. We simplified this model using stepwise model simplification (backwards and forwards) based on AIC using the function ‘step’ in R base *stats*.

In addition, we developed a structural equation model (SEM) using *iNaturalist* observations to examine relationships between moth diversity, vegetation (deciduous forest, grassland, pasture, shrub), developed land (high-, medium-, low-intensity), and developed open land cover. SEMs analyze multiple relationships at the same time while considering latent factors (i.e., unobserved relationships, such as between deciduous land cover and grassland). We specified the model to reflect the hypothesis that different components of the urban landscape (i.e., vegetation, developed land, and open space) each exert direct effects on moth diversity, and that these land cover types may covary due to the spatial structure of urban environments (e.g., lower vegetation cover in highly developed areas). Latent variables were used to group related land cover types, reducing collinearity and capturing broader ecological gradients. We fit our SEM using maximum likelihood with the ‘sem’ function in the *lavaan* R package (ver. 0.6-19) [49]. Model fit was assessed using Chi-squared (χ²), Comparative Fit Index (CFI), Tucker-Lewis Index (TLI), and Root Mean Square Error of Approximation (RMSEA). To account for shared variance between variables, we specified covariances between vegetation cover and development intensity, development intensity and developed open space, and between several vegetation variables, including between deciduous and grassland, and between grassland and shrub. We used the z-statistic to assess the significance of each path coefficient in the SEM and visualized observed relationships with the function ‘semCors’ in the R package *semPlot* [50].

We conducted two additional analyses using our on-the-ground field sampling data. We again analyzed the similarity of community composition concerning environmental similarity across sites using MRM. As with the *iNaturalist* MRM analysis, this entailed generating a distance matrix of species composition similarity across sampling sites, and for the environment: percentages of all NLCD land cover classes, as well as PM2.5, lux, temperature, humidity, and wind speed measured on the ground at time of sampling. Second, we used linear regression to analyze the relationship between moth diversity and site-level habitat features. We did not include remotely sensed land cover data in this model because the resolution of these data and our sample size resulted in low statistical power (overparameterization) and multicollinearity when these variables were included. Explanatory variables used in our model included temperature, pollution (PM 2.5), and light intensity (lux) measured during on-the-ground surveys. We simplified this model using stepwise model simplification (backwards and forwards) based on AIC using the function ‘step’ in R base *stats*. Lastly, we combined *iNaturalist* data with our on-the-ground survey data to evaluate the relationship between diversity and urbanization more broadly. Specifically, we analyzed land cover PCA, with no interactions or model simplification.

## Results

### Urban environmental variation

Land cover analyses using remotely sensed data verified that our on-the-ground sampling locations spanned a broad range of urbanization (Fig. 2a). Larger parks (Forest Park, Highland Park, Prospect Park) exhibited greater habitat heterogeneity and had more deciduous forest cover. Small parks (Maria Hernandez, Grover Cleveland, Irving Square), by contrast, had little to no deciduous tree cover and were composed almost entirely of developed land. Grover Cleveland Park stood out with only a small percentage of high-intensity developed land as it borders a private cemetery with considerable greenspace. These differences are clearly visible in satellite imagery: Forest Park is dominated by forest, Prospect Park has less tree cover, and Maria Hernandez is substantially smaller and consists mostly of developed land (Fig. 2c).

**Figure 2.**
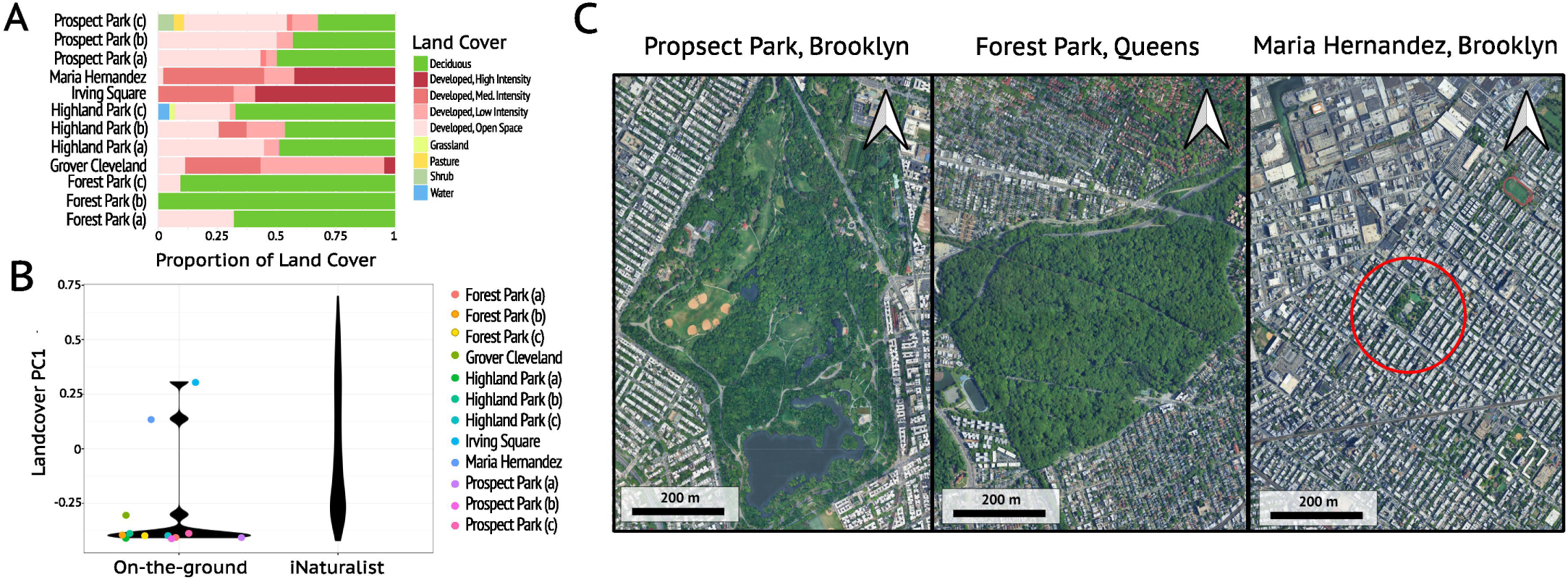
Land cover composition, principal components analysis, and site imagery of sampling locations (a) Stacked bar plot of NLCD land cover composition at on-the-ground sampling sites. Colors represent the proportion of each land cover type in 150 m buffers around each sampling location. (b) Principal components analysis of NLCD land cover for our on-the-ground sampling locations and all hex-bins used with iNaturalist data. PC1 represents high-intensity developed land cover. (c) Satellite images of sampling locations in Brooklyn and Queens, New York.

Our PCA incorporated land cover for both *iNaturalist* records and on-the-ground sites (1115 *iNaturalist* hexagonal zones, 12 on-the-ground sampling locations). The PCA captured 95.31% of the variance in the first four principal components, with PC1 capturing 44.68% of the variance. Visual inspection of the scree plot suggests that four principal components should be retained to describe variation in the dataset. The first and second principal components (PC1 and PC2) represent urbanization, with strong positive loadings for developed high-intensity land cover on PC1 (Fig. 2b) and for medium-intensity developed land cover on PC2. The third component (PC3) was negatively correlated with water and positively correlated with developed open space and deciduous tree cover. The fourth principal component (PC4) was positively correlated with developed open space and developed low-intensity land cover and negatively associated with deciduous tree cover. Grassland, pasture, and shrub land cover classifications did not load strongly on any of the first four principal components.

### Diversity of Lepidoptera

From *iNaturalist*, we extracted data from 9,377 Research Grade observations, representing 972 species, 535 genera, and 50 families (Fig. 3b). In our on-the-ground surveys, we observed a total of 69 individuals, representing 11 families, 31 genera, and 33 species (Fig. 3a). Observations were identified to species in 100% of samples. When comparing *iNaturalist* moth richness with surveyed richness, *iNaturalist* richness was markedly higher than that of our surveys for Prospect Park (110 species on *iNaturalist* versus 12 in our sampling), but similar for Grover Cleveland (two on *iNaturalist* versus three). By contrast, in two parks, we detected more species in our on-the-ground sampling: Forest Park (29 on *iNaturalist* versus 35) and Highland Park (nine on *iNaturalist* versus 19). In the two parks where no species were detected in our on-the-ground sampling (Maria Hernandez and Irving Square), there were also no observations on *iNaturalist*. Notably, we detected individuals from the family Eriocraniidae in our on-the-ground sampling, but this family was not present in the *iNaturalist* dataset. Conversely, *iNaturalist* observations documented 40 additional families not detected in our on-the-ground sampling.

**Figure 3.**
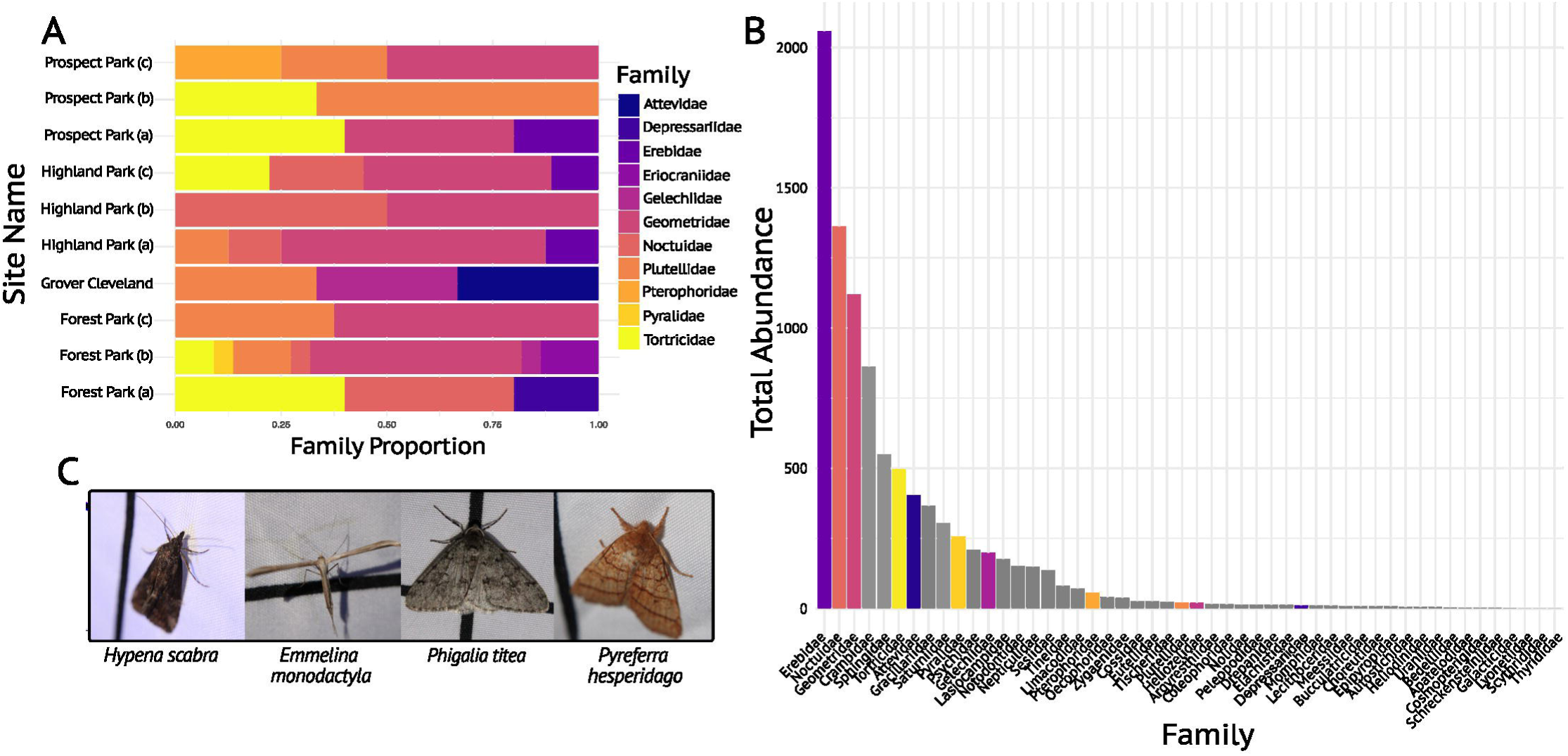
Lepidoptera family diversity from field sampling and iNaturalist data (a) Stacked bar plot of Lepidoptera Family diversity per site for on-the-ground sampling. (b) Number of observations of each Lepidoptera Family across all iNaturalist hex bins. Bars are colored by Families observed. (c) Images of moths taken during on-the-ground sampling.

Based on our on-the-ground sampling, large parks (Forest Park, Highland Park, Prospect Park) displayed the highest levels of biodiversity (1-D: 0.44–0.92, mean±sd: 0.72±0.15). Among them, Forest Park had a mean diversity index of 0.80±0.1, Highland Park had a mean diversity of 0.71±0.19, and Prospect Park had a mean diversity of 0.64±0.17. In contrast, the small parks (Maria Hernandez, Grover Cleveland, Irving Square) exhibited lower diversity (1-D: 0–0.66, mean±sd: 0.22±0.38). No species were detected at Maria Hernandez and Irving Square in our on-the-ground sampling; since Simpson’s Diversity Index is undefined in the absence of observations, we recorded 1-D as 0 for these sites.

### Community Composition

In both our *iNaturalist* data and on-the-ground sampling, locations with similar environments had similar community composition at the species level (multiple-matrix regression; on-the-ground: F=43.127, R^2^=0.501, p=0.031; *iNaturalist*: F=3892.81, R^2^=0.013, p<0.001). Species diversity is strongly negatively correlated with PC1 and PC2 (representing increased high– and medium-intensity development, respectively) across both datasets combined (PC1 estimate=-0.200±0.041, t=-4.848, p<0.001; PC2 estimate=-0.449±0.052, t=-8.692, p<0.001) (Fig. 4). Diversity was also positively correlated with PC3, representing less water and more developed open space and deciduous tree cover (estimate=0.415±0.073, t=5.654, p<0.001). Diversity was not significantly correlated with PC4 (t=0.046, p=0.963).

**Figure 4.**
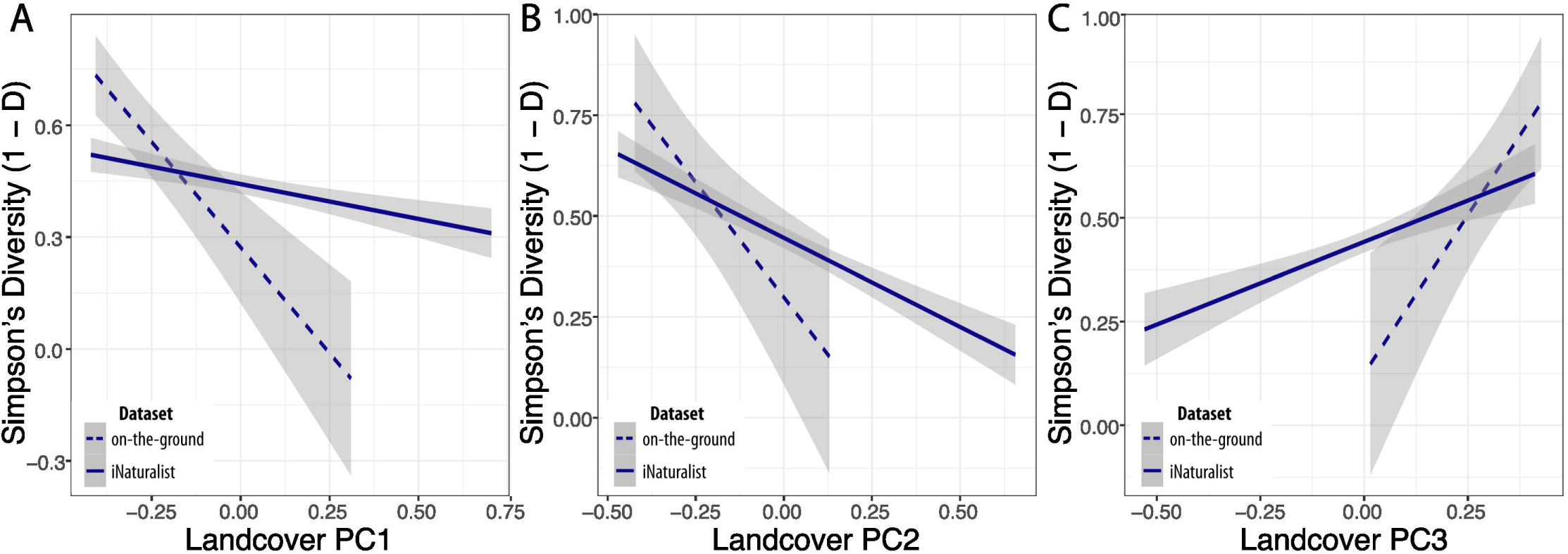
Correlations of Simpson’s Diversity Index and NLCD land cover. Simpson’s diversity (D-1) as it relates to land cover summarized by principal components analysis: (a) PC1 representing high-intensity developed land, (b) PC2 representing medium-intensity developed land, and (c) PC3 representing open developed land and deciduous tree cover.

### Environmental Correlations

Our two linear models—one using *iNaturalist* observations with remotely sensed environmental variation and the other using on-the-ground sampling with field-measured environmental variation—revealed correlational relationships between environmental features and biodiversity across both spatial scales. Our simplified linear regression of Simpson’s Diversity by land cover using *iNaturalist* and remotely sensed data retained five significant variables: medium-intensity developed land, low-intensity developed land, developed open land, shrubland, and deciduous tree cover. Diversity was negatively correlated with medium-intensity developed land cover (estimate=-0.350±0.064, t=-5.493, p<0.001) and positively correlated with deciduous tree cover (estimate=0.465±0.107, t=4.347, p<0.001) and developed open space (estimate=0.375±0.118, t=3.169, p=0.002). There was also a near-significant positive trend between diversity and low-intensity developed space (estimate=0.268±0.153, t=1.755, p=0.080), and the correlation with shrubland was non-significant (t=1.598, p=0.110). In comparison, our simplified regression of Simpson’s Diversity for on-the-ground sampling retained two variables: air pollution and light at night. Simpson’s diversity was not significantly correlated with air pollution (t=1.280, p=0.233) but was significantly negatively correlated with light at night (estimate=-0.512±0.062 lux, t=-8.214, p<0.001).

### Structural Equation Model

Our SEM exhibited an overall good fit to the data (CFI: 0.957, TLI: 0.918, RMSEA: 0.066). The chi-square statistic was significant (χ²=80.896, df=19, p<0.001), although this is expected with larger sample sizes, and the overall fit indices suggest the model effectively captured the relationships in the data.

Our SEM detected a significant positive effect of vegetation cover on diversity (estimate=1.905±0.398, z=4.791, p<0.001; Fig. 5) and a significant negative effect of developed land on diversity (estimate=-0.301±0.101, z=-2.979, p=0.003). We did not find a significant direct effect of developed open space (z=0.391, p=0.695) on diversity (Fig. 5). Significant negative covariance was observed between vegetation cover and development intensity (covariance=-0.012, z=-10.728, p<0.001), and significant positive covariance between vegetation cover and open space (covariance=0.002, z=4.191, p<0.001). Similarly, we found a significant negative covariance between development intensity and developed open space (covariance=-0.014, z=-11.707, p<0.001). We also observed weak but significant positive covariance between developed low-intensity and developed open space (covariance=0.005, z=9.382, p<0.001) and grassland and shrub (covariance<0.001, z=10.954, p<0.001). We observed weak but significant negative covariance between developed high-intensity and developed low-intensity (covariance=0.015, z=-11.237, p<0.001), and deciduous and grassland (covariance<-0.001, z=-2.650, p = 0.008). Further inspection of relationships among variables indicates that developed medium-intensity land cover is driving the direct effect of development on diversity, whereas high and low-intensity development impact diversity more indirectly via their relationships with open land and deciduous tree cover. Further, among the vegetation variables, only deciduous tree cover has a strong direct effect on diversity, whereas grassland, shrubland, and pasture all have indirect effects related to their covariance with developed land.

**Figure 5.**
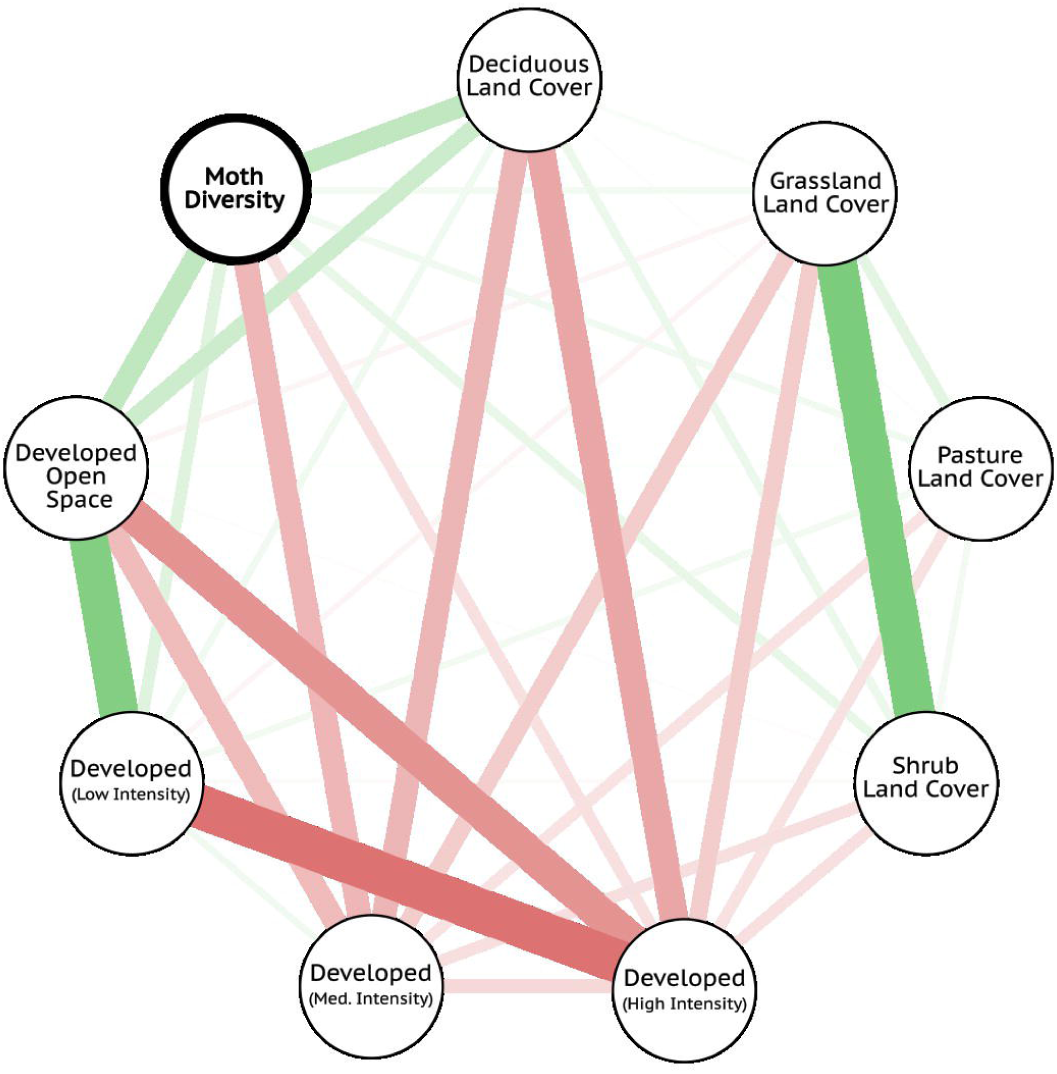
Pathways linking urban land cover to Lepidoptera diversity revealed by structural equation modeling. Relationships among predictor variables and Simpson’s diversity generated by the structural equation model. Nodes represent variables included in the model, while edges indicate associations between them. Thicker lines represent stronger relationships, with positive correlations shown in green and negative correlations in red. Edge weights are proportional to the strength of standardized regression coefficients.

## Discussion

Urbanization drastically alters landscapes, replacing naturally occurring habitats and vegetation with impervious surfaces and buildings. This process not only impacts land cover but also increases artificial light at night (ALAN), both of which can have significant consequences for biodiversity. Lepidoptera (moths and butterflies) are highly sensitive to land cover changes [31, 51], and are also influenced by habitat patch size, quality, and shape [52–53]. In this study, we examined how urbanization relates to the diversity of nocturnal Lepidoptera (moths), which perform vital ecosystem services and act as bioindicators [28]. Our findings suggest that urbanization is associated with lower moth diversity, with specific elements of the urban landscape shaping patterns of biodiversity in complex ways.

We found comparable moth community composition between locations with similar environments in both *iNaturalist* and our on-the-ground sampling datasets, and a general trend of decreasing diversity with increasing urbanization. The *iNaturalist* MRM had a low R² but was highly significant, likely reflecting high spatial coverage but heterogeneous observer effort. The on-the-ground MRM explained much more variation in community composition despite its smaller sample size and its limited spatial and temporal scope, suggesting that controlled sampling better captured ecological patterns among sites. These results suggest that moth community composition across New York City is strongly influenced by the environmental features that define different locations. Our linear models provide some insight into those features: we found strong negative correlations between moth diversity and medium-intensity urbanization, and strong positive correlations between diversity and deciduous tree cover and open land. In addition, our on-the-ground sampling found a strong negative relationship between moth diversity and light at night, with a high explained variance (R²), suggesting light at night accounts for a substantial portion of the observed variation in species diversity across sampled locations. Although several studies have found no consistent effects of light at night on biodiversity [11, 54], our results are in line with those that identify significant impacts on moth community composition [55–57]. These negative effects on biodiversity are likely attributed to how light at night disrupts moth feeding, reproduction, migration, and development [21].

Our PCAs of *iNaturalist* and on-the-ground data reinforce these findings: we found strong negative correlations between diversity and the principal components primarily defined by high– and medium-intensity development (i.e., PC1, PC2), and strong positive correlations with the principal component representing deciduous tree cover and open space (i.e., PC3). The repeated emergence of deciduous tree cover, open space, and high– and medium-intensity developed land as key variables across multiple analyses highlights the importance of these features. Our results are consistent with those of similar studies performed in metropolitan areas where highly disturbed habitats have been shown to correlate with reduced moth species richness [20, 23, 58].

While we did not perform a SEM using our on-the-ground data due to the larger sample sizes required, our SEM using *iNaturalist* data and remotely sensed land cover found, similar to our other analyses, that higher diversity is associated with greater vegetation cover and less developed land. Further inspection of these relationships revealed that although vegetation and development have direct impacts on diversity, multiple indirect effects exist. Our strongest observed path within the SEM was the direct positive effect of vegetation on diversity while development showed a weaker but significant negative direct effect. Indeed, the strong negative covariance between vegetation and development and a strong positive covariance between vegetation and developed open land, suggests that characteristics of more intense urbanization may indirectly drive reduced moth diversity by altering land cover compositions. Taken together, these results indicate that high– and medium-intensity development reduce moth diversity not only directly, but also indirectly through their association with reduced vegetation cover. Together, these results suggest that developed land conversion drives species declines independently and through strong indirect effects. This finding points to potential avenues for moth conservation through the maintenance of tree cover, vegetation, and open space even in highly developed parts of the city. Therefore, urban planning efforts focused on restoring parks and increasing vegetation across the city may make habitat more suitable for local moth populations.

Surprisingly, we did not find a relationship between air pollution and moth diversity at our on-the-ground sampling study sites. This finding contradicts our expectations, as researchers have documented declines across various taxa in response to higher levels of air pollution [59–60]. However, others suggest that the negative effects of air pollution on insects may have been overestimated [61]. It is also possible that our findings were influenced by confounding variables such as temperature and wind speed, as higher levels of particulate matter (PM2.5) coincided with warmer temperatures and lower wind speeds during sampling. A longer temporal scale may be required to detect a correlation between air pollution and moth biodiversity, given the high variation and sensitive nature of this variable. These factors highlight the complexity of isolating individual environmental drivers of diversity decline and underscore several limitations of our study.

We also acknowledge the inherent limitations of using *iNaturalist* data, which can be influenced by biases in socioeconomic factors and uneven sampling efforts. Research suggests that areas with higher socioeconomic status often experience greater sampling efforts by community scientists [62]. Taxonomic biases also exist; charismatic fauna, such as butterflies, are reported more frequently despite the greater overall diversity of moth species [63]. For these reasons we conducted on the ground surveys to reinforce our *iNaturalist* analyses. While our field data overall reiterated the same findings, it also further highlighted the limitations of citizen science data. For example, our field sampling documented high diversity in Forest and Highland Parks, even though these parks have few moth records on *iNaturalist*. In fact, our on-the-ground sampling documented more moth species over a single evening at three locations in each of these parks than have ever been documented on *iNaturalist* for the entire park. Forest Park had 29 records on *iNaturalist* in over 20 years of observations in New York City, versus our 35 observations from our on-the-ground surveys; Highland Park is similar with 9 *iNaturalist* observations versus our 19. Indeed, we detected members of a family (Eriocraniidae) that had never been observed in New York City on *iNaturalist*, perhaps because of their particularly small size (∼10 mm wingspan) and early spring emergence. However, the *iNaturalist* data recorded 40 families not observed in person. Moreover, *iNaturalist* has many records of larval stage moths, which would not be recorded during a light trap survey, and other adult stages of certain species which are difficult to record via light trapping due to variation in phototaxis [41]. The discrepancies between datasets highlight the importance of integrating *iNaturalist* data with on-the-ground field surveys to develop a more comprehensive understanding of biodiversity in a given area, and particularly in cities where habitat is highly heterogeneous and strong sampling biases can arise.

In conclusion, our analyses revealed significant relationships between moth diversity and urbanization at both landscape and local spatial scales. Our landscape-level analysis found that higher vegetation cover and developed open space support greater moth diversity, while our on-the-ground sampling revealed negative effects of light at night. Further, our findings highlight the importance of using advanced modeling approaches, such as SEM, alongside linear regression to reveal hidden or unobserved relationships in environmental data. Understanding these relationships is essential for designing targeted conservation strategies, as singular actions such as increasing vegetation alone, may be insufficient. We hope this study sparks further research into urban moth diversity, particularly given the essential role these organisms serve in the world’s ecosystems.

## Acknowledgements

We are thankful to Colin Ainsworth and Cassie Lustig who assisted in field-based data collection. We would also like to recognize New York University’s DURF and Undergraduate Research Assistant Programs for making this research possible.

## Data Availability

All data and associated code are archived on Zenodo with doi: 10.5281/zenodo.14756963

